## Supplementary Figures for "Immunometabolic determinants of long-term response in leukemia patients receiving CD19 CAR T cell therapy"

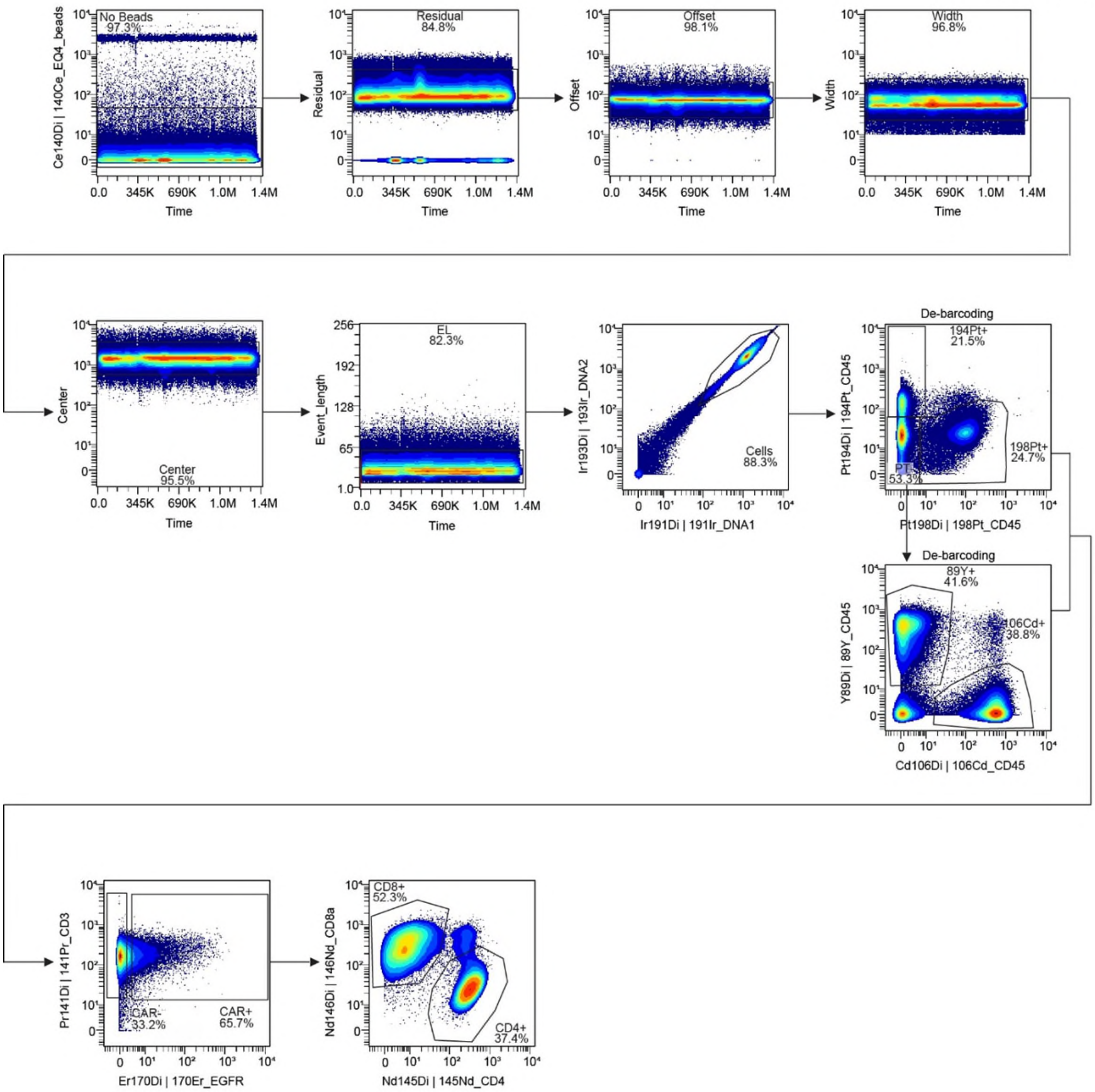


**Supplementary Fig. 1: Representative of gating strategy to detect CAR- and CAR+ T cells by mass cytometry.** CyTOF data were cleaned using the Gaussian parameter discrimination method, and live cells were detected by excluding beads (140Ce) and manually gating on DNA (191Ir/193Ir) and viability (195Pt). Patients' samples were manually de-barcoded (CD45; 194Pt/198Pt/89Y/106Cd), and CAR T and non-CAR T cells were identified using anti-CD3 (141Di) and anti-huEGFRt (170Er), followed by CD4 (145Nd) and CD8 (146Nd) metal-tagged antibodies.

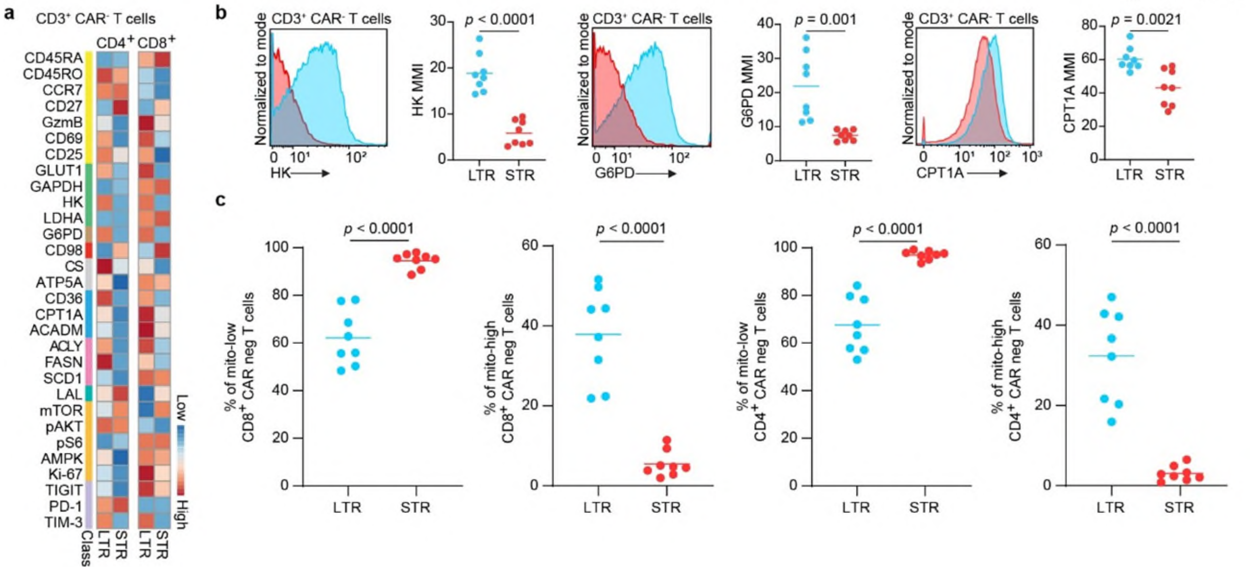


**Supplementary Fig. 2: CD3+ CAR- T cells of long-term responders show increased mitochondrial mass and expression of key enzymes involved in oxidative phosphorylation, fatty acid oxidation, and pentose phosphate pathway activity. a** Heat map indicating z-score normalized median expression of all CyTOF markers assessed in CD4+ and CD8+ CAR- T cells. **b** Representative histograms and analysis of statistically significant proteins in CD3+ CAR- T cells. **c** Analysis of MitoTracker expression in CD4+ and CD8+ CAR- T cells products.

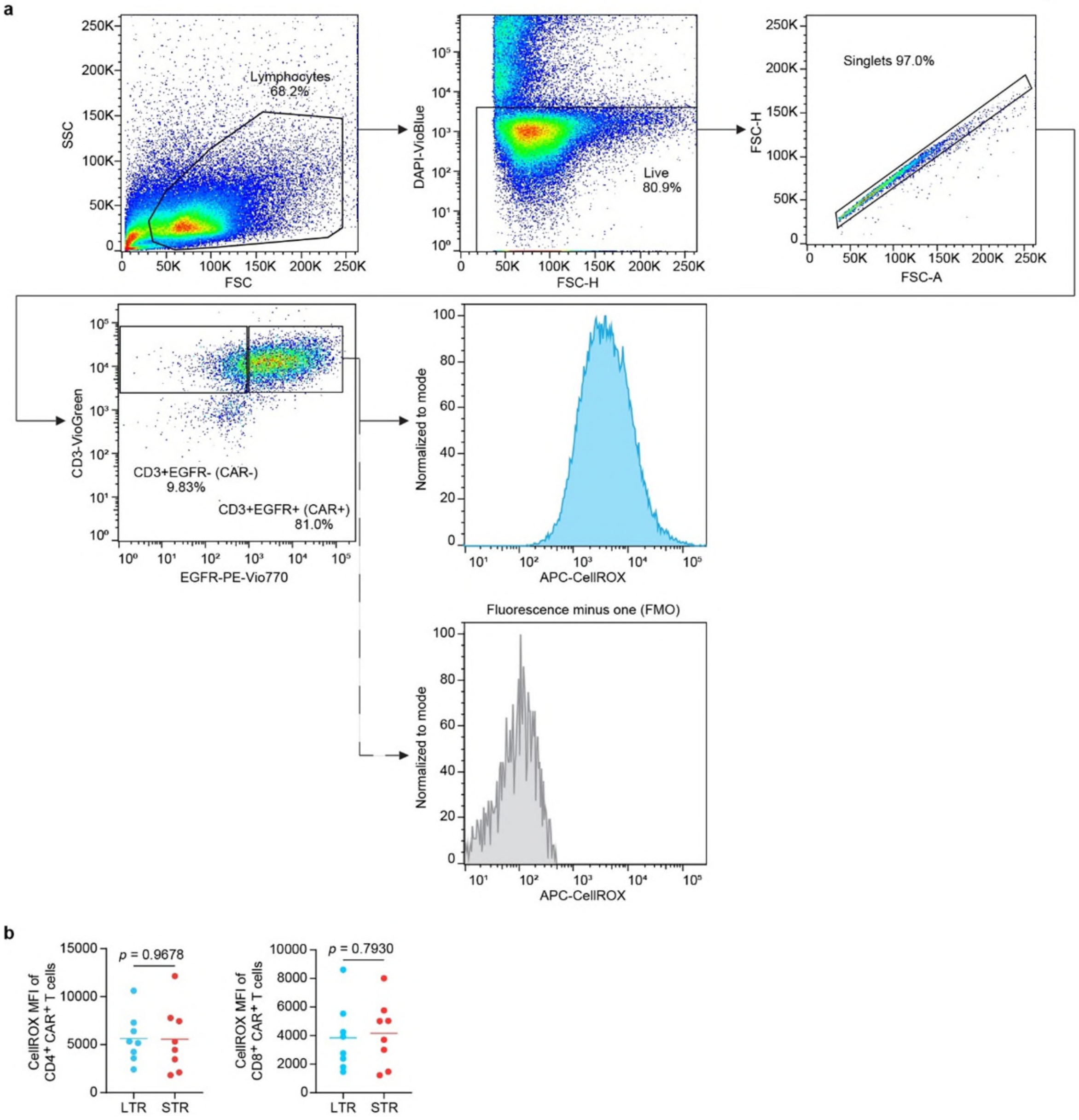


**Supplementary Fig. 3: Reactive oxygen species assessment by flow cytometry. a** Representative of gating strategy. All samples were FSC and SSC gated. DAPI negative cells were then gated, followed by FSC-A/FSC-H gating to select singlet cells. CAR T cells were detected using CD3 and EGFR staining, and reactive oxygen species level was assessed by measuring the CellROX fluorescent probe expression (%). **b** Analysis of CellROX expression in CD4+ and CD8+ CAR+ T cells products.

Supplementary Fig. 4

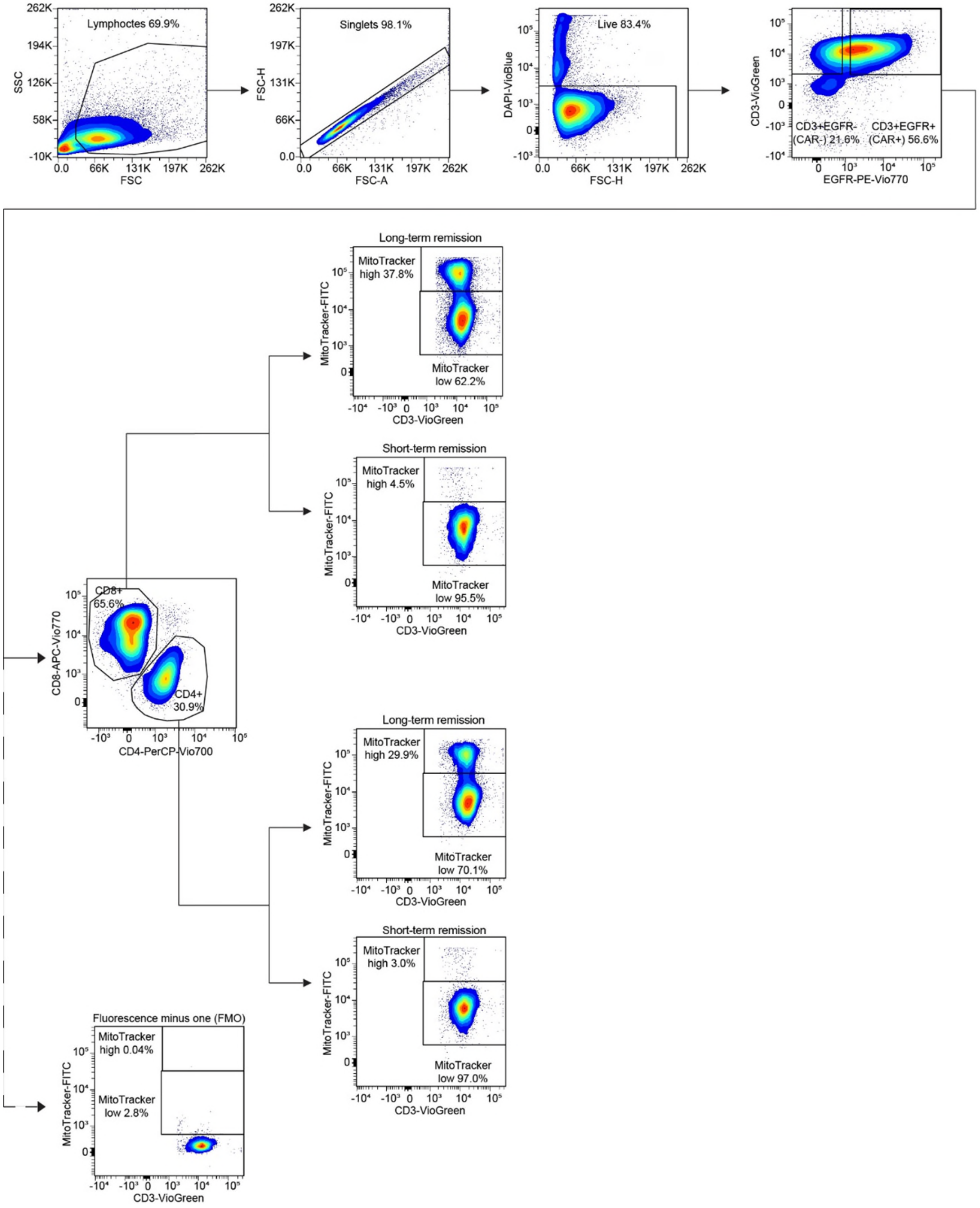

**Supplementary Fig. 4: Mitochondrial mass assessment by flow cytometry.** Representative of gating strategy. All samples were FSC and SSC gated, followed by FSC-A/FSC-H gating to select singlet cells. DAPI negative cells were then gated, and CAR and non-CAR T cells were detected using CD3 and EGFR staining. Mitochondrial mass was assessed by measuring the MitoTracker fluorescent probe expression (%).

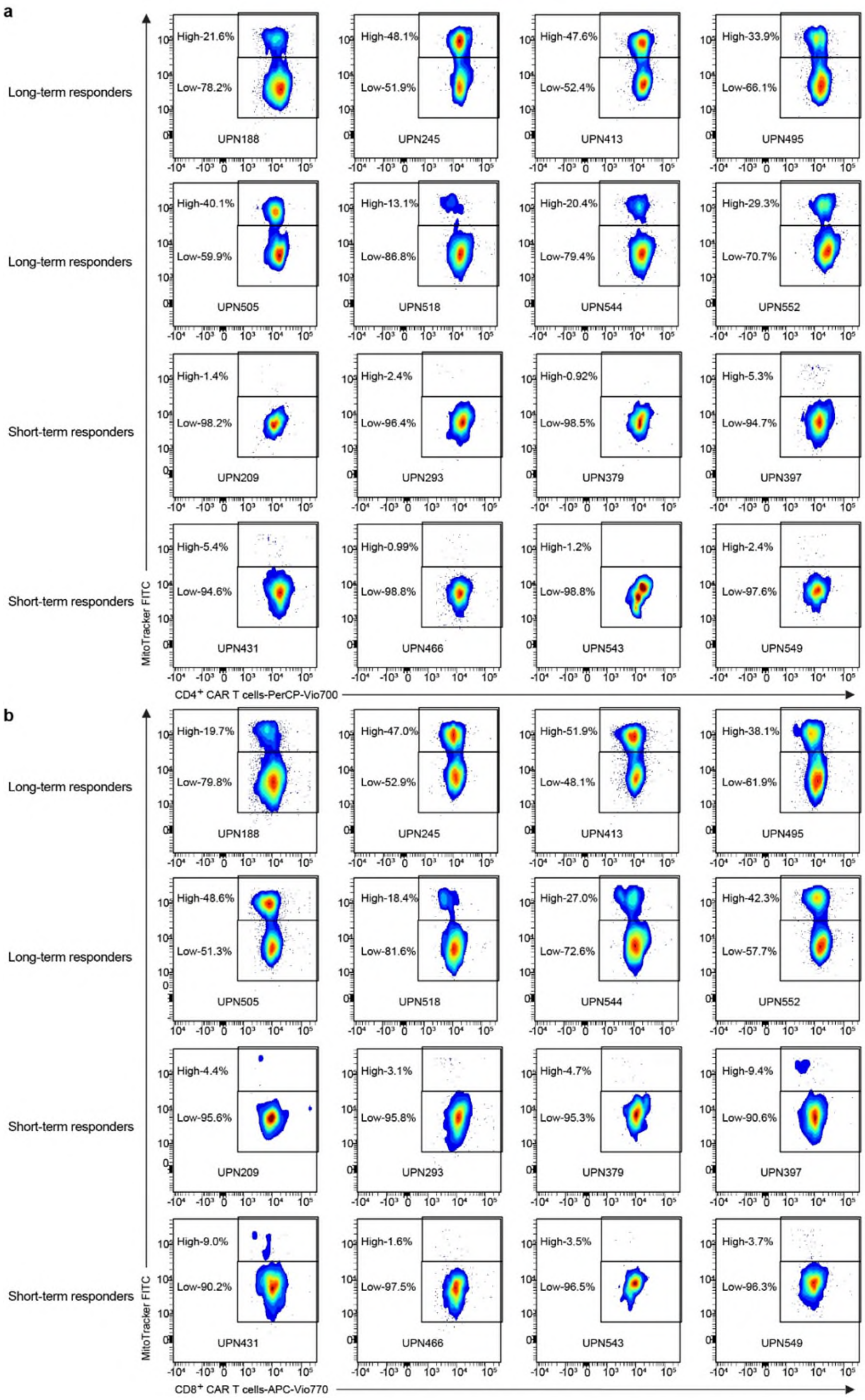
a
Long-term responders
High-21.6%
Low-78.2%
UPN188
High-48.1%
Low-51.9%
UPN245
High-47.6%
Low-52.4%
UPN413
High-33.9%
Low-66.1%
UPN495
Long-term responders
High-40.1%
Low-59.9%
UPN505
High-13.1%
Low-86.8%
UPN518
High-20.4%
Low-79.4%
UPN544
High-29.3%
Low-70.7%
UPN552
Short-term responders
High-1.4%
Low-98.2%
UPN209
High-2.4%
Low-96.4%
UPN293
High-0.92%
Low-98.5%
UPN379
High-5.3%
Low-94.7%
UPN397
Short-term responders
High-5.4%
Low-94.6%
UPN431
High-0.99%
Low-98.8%
UPN466
High-1.2%
Low-98.8%
UPN543
High-2.4%
Low-97.6%
UPN549
MitoTracker FITC
CD4+ CAR T cells-PerCP-Vio700
b
Long-term responders
High-19.7%
Low-79.8%
UPN188
High-47.0%
Low-52.9%
UPN245
High-51.9%
Low-48.1%
UPN413
High-38.1%
Low-61.9%
UPN495
Long-term responders
High-48.6%
Low-51.3%
UPN505
High-18.4%
Low-81.6%
UPN518
High-27.0%
Low-72.6%
UPN544
High-42.3%
Low-57.7%
UPN552
Short-term responders
High-4.4%
Low-95.6%
UPN209
High-3.1%
Low-95.8%
UPN293
High-4.7%
Low-95.3%
UPN379
High-9.4%
Low-90.6%
UPN397
Short-term responders
High-9.0%
Low-90.2%
UPN431
High-1.6%
Low-97.5%
UPN466
High-3.5%
Low-96.5%
UPN543
High-3.7%
Low-96.3%
UPN549
MitoTracker FITC
CD8+ CAR T cells-APC-Vio770

**Supplementary Fig. 5: Mitochondrial mass assessment of pre-infusion CAR T cells in products of short- and long-term responders.** Biaxial plots showing Mitotracker expression level (%) in CD4+ (**a**) CD8+ **(b)** CAR+ T cells.

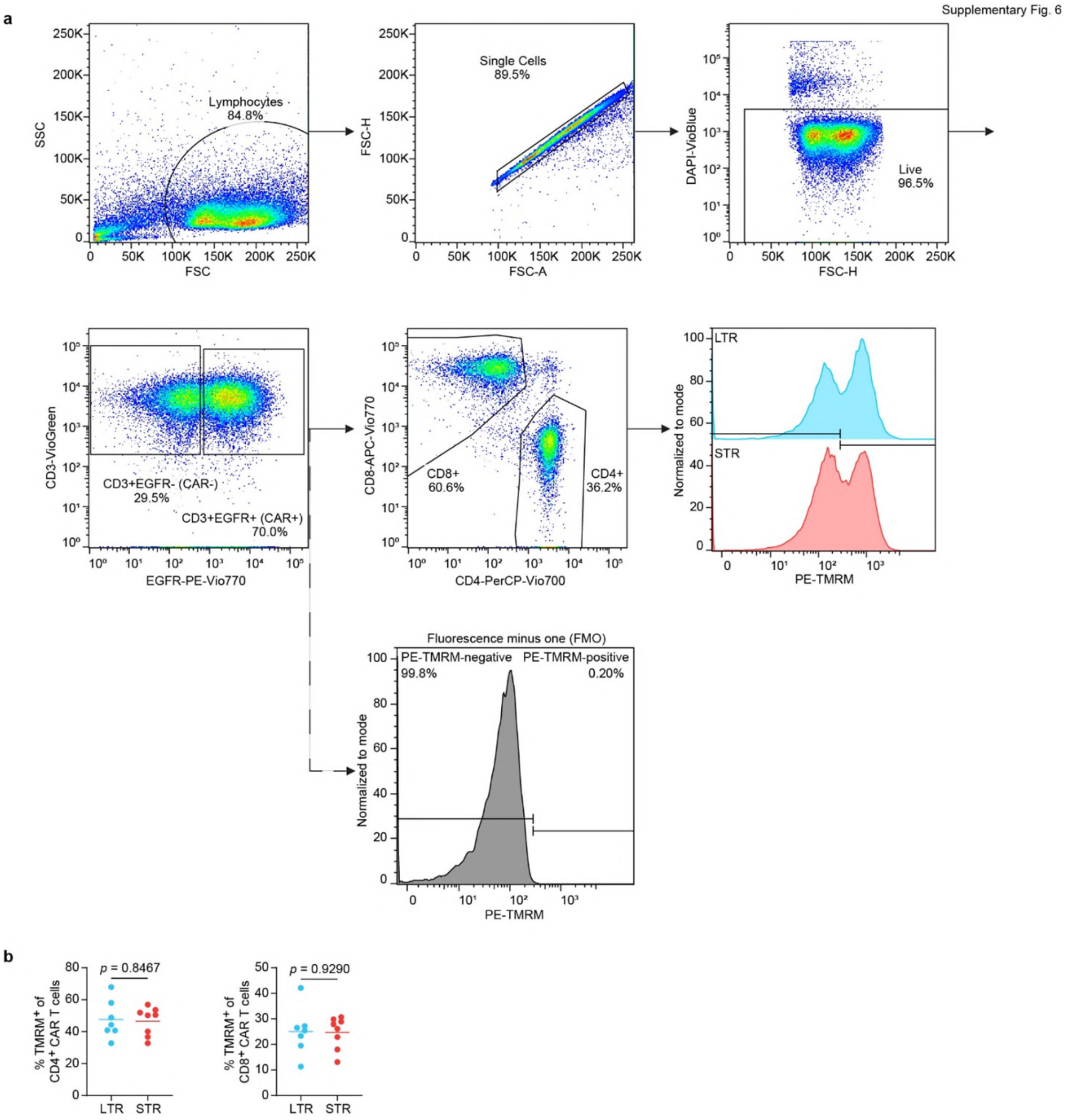


**Supplementary Fig. 6: Mitochondrial membrane potential assessment by flow cytometry.** **a** Representative of gating strategy. All samples were FSC and SSC gated, followed by FSC-A/FSC-H gating to select singlet cells. DAPI negative cells were then gated, and CAR cells were detected using CD3 and EGFR staining. Mitochondrial membrane potential was assessed by measuring the TMRM (Tetramethylrhodamine methyl ester) dye expression (%). **b** Analysis of TMRM expression in CD4+ and CD8+ CAR+ T cells pre-infusion products.

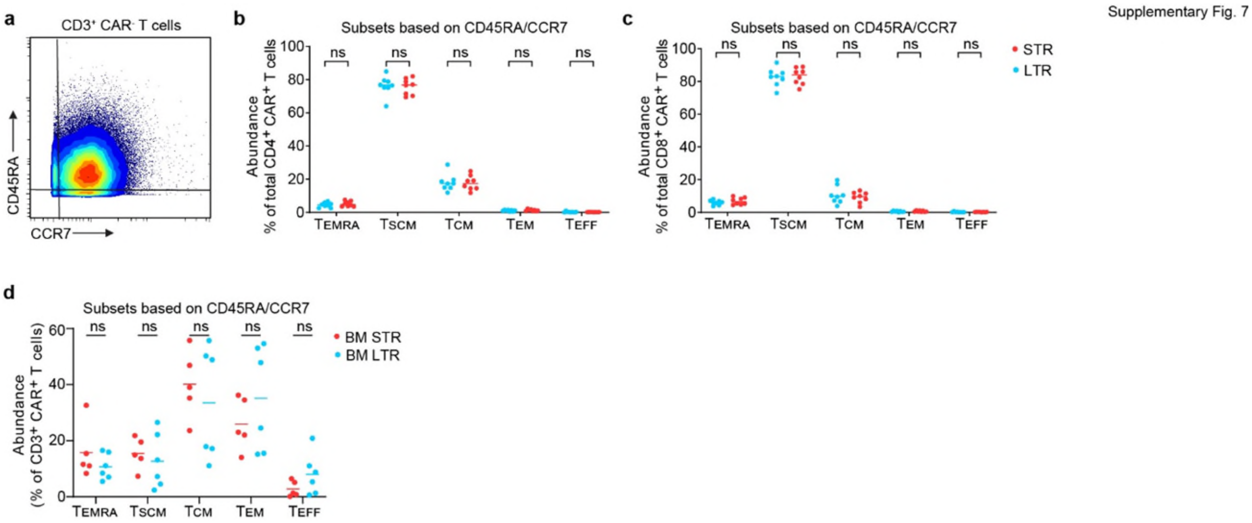


**Supplementary Fig. 7: T cell differentiation subsets abundance by flow cytometry. a** Representative biaxial plot to measure CAR T cell subsets using CD45RA and CCR7. Abundance of canonical T-cell subsets in CD4+ (**b**) and CD8+ (**c**) CAR+ T cells of pre-infusion products of short- and long-term responders. **d** Abundance of canonical T-cell subsets in CD3+ CAR+ T cells of day 28 post-infusion bone marrow samples of short- and long-term responders.
